# Tree of Mutually Exclusive Oncogenes

**DOI:** 10.1101/2022.02.07.479425

**Authors:** Mohammadreza Mohaghegh Neyshabouri, Jens Lagergren

## Abstract

Identifying cancer driver genes and their interrelations is critical in understanding cancer progression mechanisms. In this paper, we introduce ToMExO, a probabilistic method to infer cancer driver genes and how they affect each other, using cross-sectional data from cohorts of tumors. We model cancer progression dynamics using a tree with sets of driver genes in the nodes. This model explains the temporal orders among driver mutations and their mutual exclusivity patterns. We introduce a dynamic programming procedure for the likelihood calculation and build an MCMC inference algorithm. Together with our engineered MCMC moves, our efficient likelihood calculations enable us to work with datasets having hundreds of genes and thousands of tumors in the datasets. We demonstrate our method’s performance on several synthetic datasets covering various scenarios for cancer progression dynamics. We then present the analyses of several biological datasets using the ToMExO method and validate the results using a set of method-independent metrics.

## Introduction

Cancer is a disease caused by evolutionary processes involving the accumulation of somatic mutations in the genome. Some mutations in so-called cancer driver genes confer selective advantages to the cells harboring them, resulting in cancer progression. These mutations are called driver mutations. Identifying the critical driver genes and the interplay between their mutations is crucial for a broad set of research and clinical applications, including choosing targets for new drugs, the prognosis of individual tumors, and designing patient-specific treatment plans. In abundance of so-called passenger (or background) mutations, with no decisive effect on the tumor evolution, we need to study cross-sectional data from cohorts of tumors to be able to identify the driver genes and the way their mutations affect each other.

Cancer progression has been extensively studied through an evolving range of models during recent years. Several models including oncogenetic trees [1], mixture of oncogenetic trees [2], conjunctive Bayesian networks [3] and hidden-variable oncogenetic trees [4] are designed to model the temporal order of individual driver mutations. Some other studies have tried to identify patterns of mutual exclusivity among specific driver mutations. The driver mutations typically affect the tumor evolution through disruptions in some biological pathways or protein complex structures. Therefore, the selective advantage provided by one driver mutation may exhaust that of another, leading to mutual exclusivity patterns among such mutations. Moreover, a set of mutations might be so fatal that one does not allow enough time for any of the rest to occur, leading to mutual exclusivity patterns in a different way. TiMEx [5] can be used to identify modules of mutually exclusive driver mutations. Several papers have studied the simultaneous identification of mutually exclusive sets of mutations and their temporal order. Pathway linear progression model [6] tries to find mutually exclusive sets and order them in a linear structure. A probabilistic solution for training such a model is later introduced in [7]. The TiMEx model was also later extended to pathTiMEx [8], which can be used to identify a partially ordered set of mutually exclusive modules. Several more complex models, including network aberration model [9] and mutual hazard networks [10] have also been proposed. Instead of finding sets of related mutations or a temporal order for them, these models can be used to capture the effects of all driver mutations on all other driver mutations. However, they suffer from severe computational complexity and over-fitting issues, limiting their application to datasets with very few genes.

In this paper, we introduce a probabilistic progression model for cancer. Given a dataset including information on the presence/absence of mutations in a set of genes in a cohort of tumors, we identify the cancer driver genes, group them as sets of genes showing mutual exclusivity patterns in their mutations, and arrange these sets of genes in a tree structure representing the order in which they get mutated and push the cancer forward. We introduce a computationally efficient dynamic programming procedure for likelihood calculations, used as the core of a Markov Chain Monte Carlo (MCMC) algorithm to make inference based on our model. We have also designed a set of novel structural moves, enabling us to explore our model space efficiently. This work improves the algorithm introduced in [7] in several ways. Firstly, the modeling power is improved from a single linear chain to a tree-like structure. Furthermore, the model selection issues are resolved by introducing flexible MCMC moves that split and merge nodes in various ways. Similar to the model used in [7], our model in this paper allows for false positive and false negative errors in the dataset. Although the parameters for the probability of each of these error types can be sampled in the MCMC iterations (the way [7] does it), we introduce a dynamic programming procedure to find the minimum possible values for these parameters, given a fixed structure. We use this procedure inside the MCMC iterations to speed up the training algorithm. After finding the proper structure, we fine-tune the parameters using gradient descent. In this way, we gain a significantly improved computational complexity compared to the method in [7]. We call our method ToMExO, standing for “Tree of Mutually Exclusive Oncogenes”. In this paper, we also apply ToMExO to analyze several biological datasets, including uterine corpus endometrial carcinoma, cholangiocarcinoma, and glioblastoma multiforme. Our results provide some new insights into the studied cancer types.

The paper is organized into three main parts as follows. Section 1 describes the method, where we start with introducing our cancer progression model in section 1.1. We continue with describing the probabilistic model for our data generation process in section 1.2 and introducing our likelihood calculation algorithm in section 1.3. We finish the method section by going over our inference algorithm in section 1.4. In section 2 we demonstrate our performance in a set of synthetic data simulations. Finally, in section 3, we present our analyses of several biological datasets and validate our results using two method-independent measures.

## 1 Method

### 1.1 Cancer progression model

We model cancer progression using a driver tree (*V, E*). Except for the root, the nodes include non-overlapping non-empty subsets of the driver genes. We denote the set of genes in node *v* ∈ *V* by *D*_*v*_. Each edge (*u, v*) ∈ *E* has a *firing* probability denoted by *f*_*v*_ ∈ (0, 1]. Fig. 1-A shows an example of a progression model.

**Fig 1.**
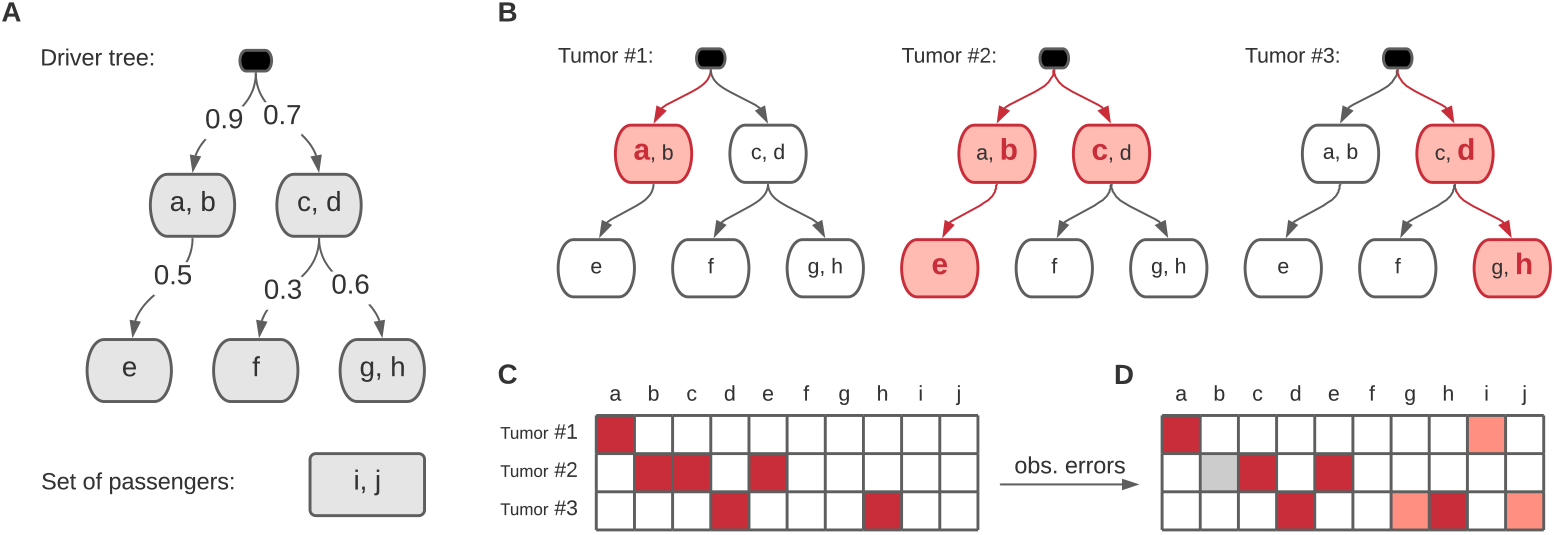
**A**. An example cancer progression model. **B**. Three tumors evolved following the model. The *firing* edges are shown in red. The mutated nodes are filled in red and the genes with driver mutations are shown in red bold font. **C**. Representation of the driver mutations in the example tumors using a binary matrix. The blocks filled in red represent the 1’s of the binary matrix. **D**. The observed dataset including errors in forms of false positives (light red) and false negatives (gray).

The model explains the tumor progressions as follows. Starting from the root, each node has a chance to get *mutated* according to its firing probability. All the mutated nodes, get a driver mutation in one and only one of their genes, chosen uniformly in random. The children of the mutated nodes may get mutated according to their firing probabilities and the cancer progresses this way. For example, consider the model shown in Fig. 1-A. Three tumors evolved according to this model are shown in Fig. 1-B. In these example tumors, the mutated nodes are colored in red and the mutated driver genes are shown using bold red font.

Given a list of mutations identified in a set of tumors, we can represent the data using a binary matrix with the tumors in the rows and the genes in the columns. Each element (*m, n*) is set to one if tumor *m* has a mutation in gene *n*. Following our example scenario in Fig. 1, the tumors shown in Fig. 1-B would result in the matrix shown in Fig. 1-C, if we had all the drivers, and no background mutations in our dataset. However, the observed data generally include a large number of background mutations. Such mutations can happen in driver genes as well. For instance, the selective advantage of mutation in a specific driver gene, may have been exhausted by an earlier mutation in another driver gene of the same protein complex. The background mutations, together with technical errors resulting in having non-real mutations in the data can be collectively seen as *false positives* in our data. On the other hand, some driver mutations may be lost due to technical issues such as low coverage of the reads covering those mutations. This can lead to what we call *false negatives* in our data.

Considering the complete list of genes with driver mutations as a perfect dataset, the aforementioned false positives and false negatives can be seen as *errors* in the data. In our example scenario, such errors lead to the observed matrix shown in Fig. 1-D.

### 1.2 Generative process

Let *B* be our binary matrix of shape *M* × *N*, including information on *N potentially* driver genes in *M* tumors. The element in the *m*^th^ row and the *n*^th^ column of the matrix, denoted by *B*_*m,n*_, is equal to one if and only if the *m*^th^ tumor has at list one mutation in the *n*^th^ gene. We denote the *m*^th^ row and the *n*^th^ column of the matrix by *B*_*m*,:_ and *B*_:,*n*_, respectively.

We call the genes not placed into the driver tree as passenger genes and denote the set of passenger genes by *P*. We denote our progression model by *T* = (*V, E*, {*f*_*v*_}_*v*∈*V*_, {*D*_*v*_} _*v*∈*V*_, *P*). Note that *T* including all the information about the topology of the tree, alongside the genes and the firing probabilities assigned to its vertices, as well as the set of passengers.

A progression model *T*, together with false positive probability *ϵ* and false negative probability *δ* form a probabilistic generative model with a graphical model shown in Fig. 2-A. The matrix *B*\* is the latent dataset, where 1’s represent the genes with driver mutations. Adding false positives and false negatives to *B*\* results in the observed matrix *B*. The likelihood of our observed data can be written as

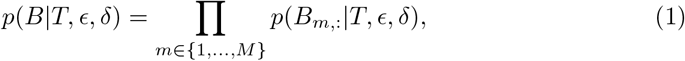

where

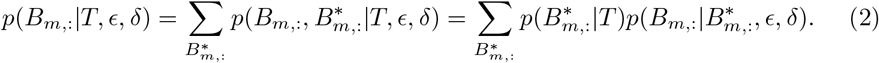

While the summation is to be taken over all 2^*N*^ states of 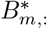, not all of these states have non-zero 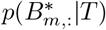, and the likelihood can be calculated in an efficient way introduced in the following section.

**Fig 2.**
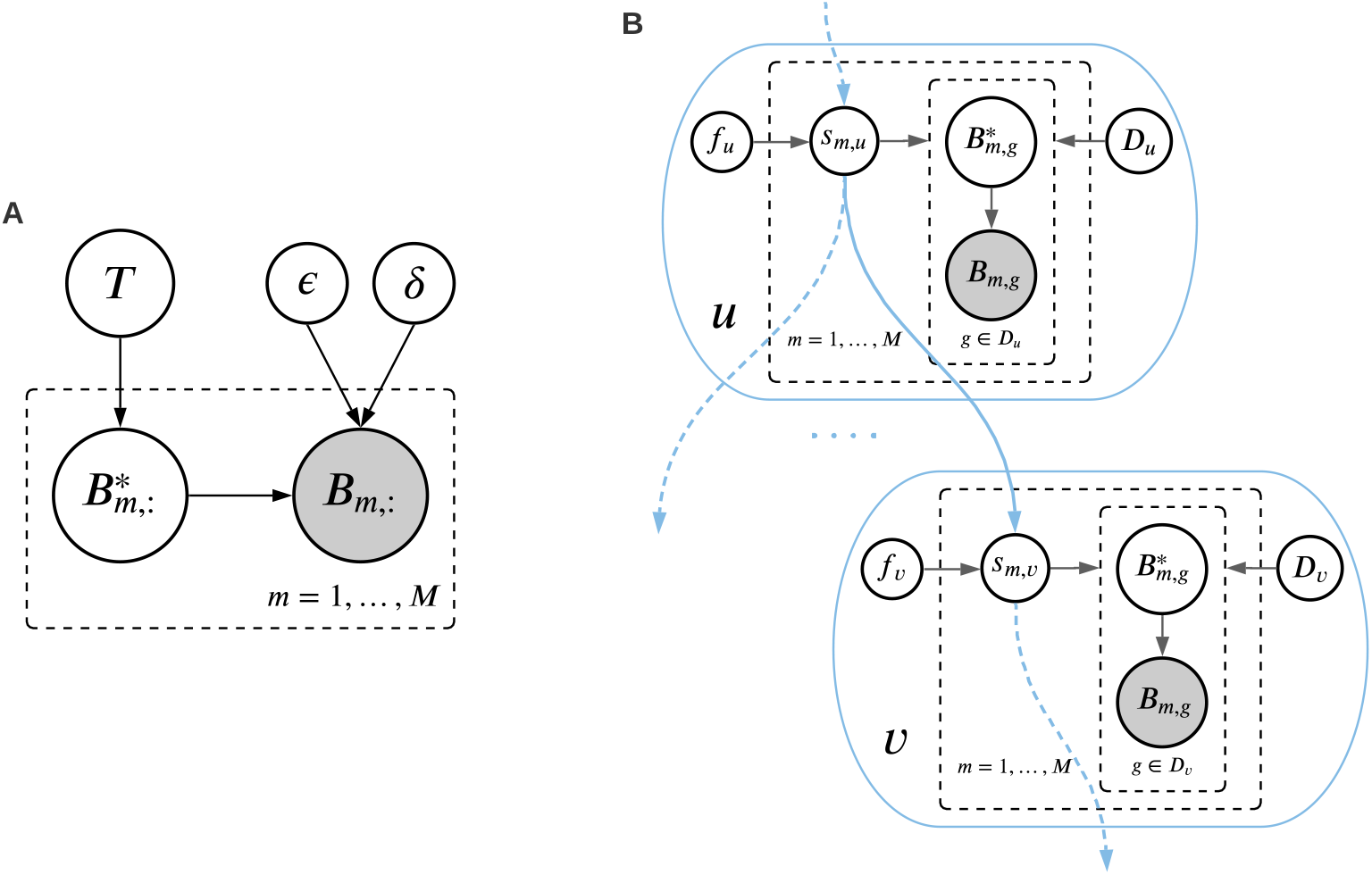
**A**. Graphical model of the generative process, where *T* is the progression model and *ϵ* and *δ* are the probabilities of false positive and false negative, respectively. The latent and observed mutation matrices are denoted by *B*\* and *B*, respectively. **B**. Graphical model of the variables in the nodes of the driver tree. The binary variable *s*_*m,u*_ is equal to one if and only if node *u* is mutated in tumor *m*.

### 1.3 Dynamic Programming for likelihood calculations

We use a dynamic programming algorithm over the tree to calculate the dataset likelihood in a computationally efficient manner. The graphical model in Fig. 2-B shows the dependencies between the variables inside the nodes of a driver tree. In this figure, the *s* variables are binary variables showing the state of nodes in the tumors (healthy/mutated). We have:

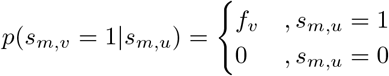

For a set of gene indices *A*, we use the notation *B*_*m,A*_ to refer to the elements of *B* in the *m*^th^ row and columns in *A*. Denoting the number of ones and zeros observed in 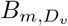 by *o*_*m,v*_ and *z*_*m,v*_, respectively, we define the variables Λ_*m,v*_ and Γ_*m,v*_ as follows:

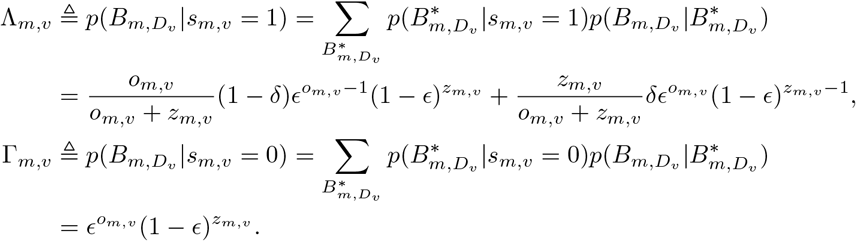

To set up our dynamic programming variables, we define *D*_*v*↓_ as the set of gene indices corresponding to vertex *v* or its descendants. We define the dynamic programming variables Ψ_*m,v*_ and Ω_*m,v*_ as:

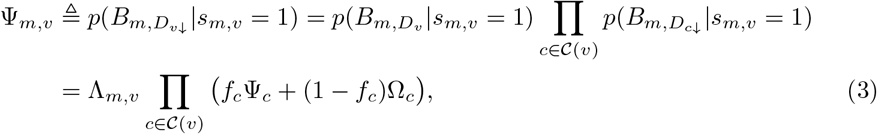

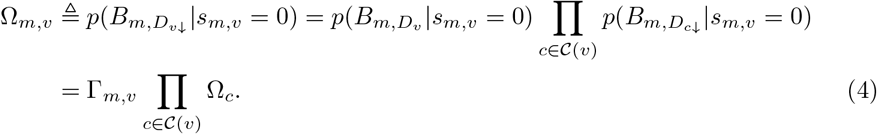

Calculating the Ψ and Ω variables following a post-order traversal on the tree, we get

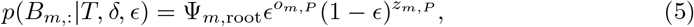

where *o*_*m,P*_ and *z*_*m,P*_ are the number of ones and zeros observed in *B*_*m,P*_, respectively.

### 1.4 Inference Algorithm

In this section, we introduce our Markov Chain Monte Carlo algorithm for making inferences using our model. Given a dataset *B*, the objective is to find a maximum likelihood estimate of the model *T* and error parameters *ϵ* and *δ*. In this paper, we restrict ourselves to models *T* = (*V, E*, {*f*_*v*_}_*v*∈*V*_, {*D*_*v*_}_*v*∈*V*_, *P*) with firing probabilities {*f*_*v*_}_*v*∈*V*_ matching the *empirical estimations*, as explained in the following. Let the topology (*V, E*) and the gene assignments ({*D*_*v*_}_*v*∈*V*_ and *P*) be given. In order to estimate the firing probability of an edge (*u, v*), denoted by *f*_*v*_, we denote the number of tumors with an observed mutation in *u* by 𝒳_*B,u*_. Similarly, we denote the number of tumors with observed mutations in both *v* and its parent *u* by 𝒴_*B,v*_. We set *f*_*v*_ = 𝒴_*B,v*_*/* 𝒳_*B,u*_ as an estimate of the firing probability, assuming few false positives and false negatives in the data.

We further restrict ourselves to error parameters *ϵ* and *δ* tuned to the model *T*. To this end, we use a dynamic programming procedure to calculate the minimum number of false positive and false negative events in the data, given a fixed progression structure. In this way, we can calculate an *empirical estimation* of the parameters *ϵ* and *δ* with a single post-order traversal of the driver tree. Our empirical error estimation algorithm is described in details in S1 Text.

We initialize our MCMC sampler to a star tree, with each gene having its own node in the driver tree. We call such single-gene nodes in the first layer of the driver tree as *simple nodes*. Following a Metropolis-Hasting framework, we use a set of possible structural moves to propose new candidate trees. The proposed trees may then get accepted based on their Metropolis-Hasting acceptance ratio. In order to explore the space of progression models in an efficient way, we have designed several types of structural moves, including various topological moves, as well as gene assignment modifications. Fig. 3 shows a few example moves. Our structural moves include:

- “Vertical merge”, merging a leaf node into its parent (Fig. 3-B), and its reverse move called “vertical split”,
- “Horizontal merge”, merging two sibling leaves (Fig. 3-C), and its reverse move called “horizontal split”,
- “Attach from passengers”, attaching a new node containing a subset of passenger genes (Fig. 3-D,), and its reverse move called “detach into passengers”,
- “Attach from simple nodes”, attaching a new node containing the genes from a subset of simple nodes (Fig. 3-E,), and its reverse move called “detach into simple nodes”,
- “P2D gene move”, moving a single gene from the set of passengers to move into an existing node (Fig. 3-F), and its reverse move called “D2P gene move”,
- “S2D gene move”, moving the gene in a simple node into an existing node (Fig. 3-G), and its reverse move called “D2S gene move”,
- “Gene swap”, swapping the genes between a node and its parent (Fig. 3-H),
- “SPR”, subtree pruning and regrafting to modify the tree structure (Fig. 3-I).

**Fig 3.**
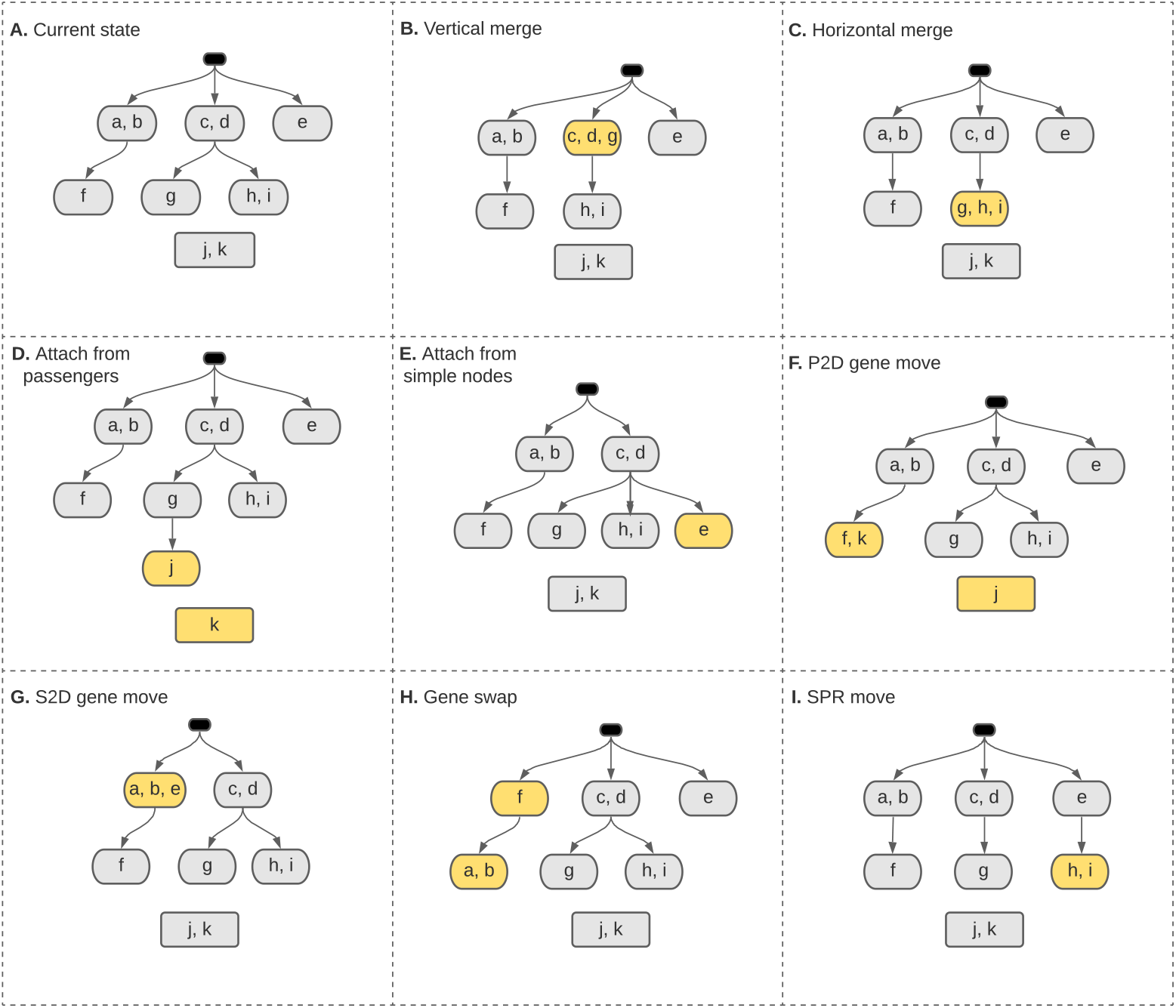
Different types of structural moves used in our MCMC inference algorithm.

After generating a set of samples, we select the sample with maximum likelihood as the output progression model and fine-tune the error parameters *ϵ* and *δ* using gradient descent with a few iterations over the selected driver tree. A pseudo-code of the inference algorithm with detailed explanations are included in S1 Text.

## 2 Synthetic Data Experiments

In this section, we use synthetic data simulations to demonstrate the efficiency of our inference algorithm in the scenarios with generative progression models shown in Fig. 4. For each scenario, we constructed 16 datasets, with equal false positive and false negative probabilities in {0.001, 0.01, 0.05, 0.1} and the number of tumors in {50, 100, 200, 500}. We used our inference algorithm with 10 MCMC chains and 100*k* iterations throughout our synthetic data experiments.

**Fig 4.**
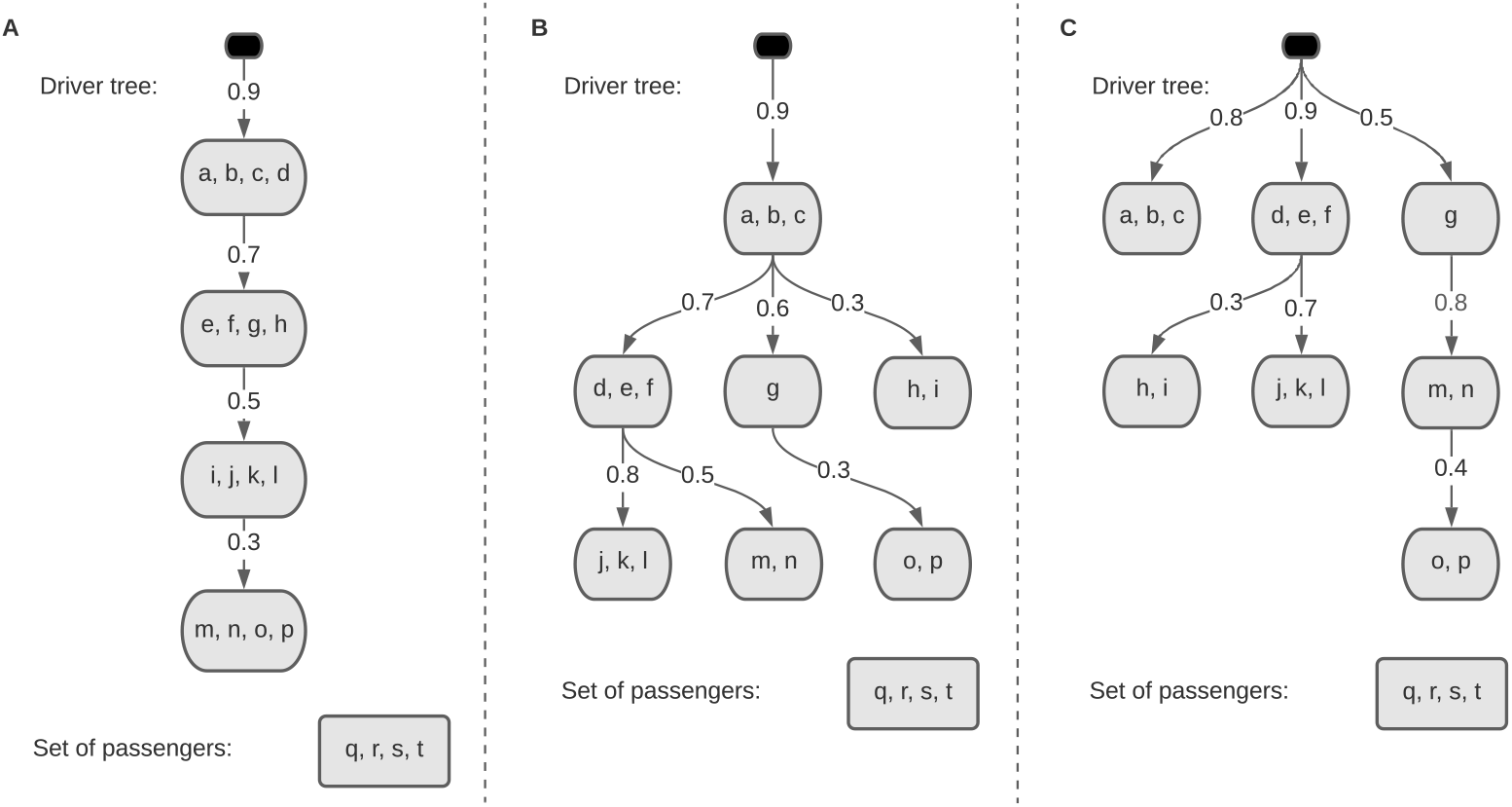
Generative progression model for the synthetic data experiments with **A**. Linear model **B**. Tree-like model **C**. Mixture of trees

Each progression model *T* implies a set of mutual exclusivity and progression relations among pairs of genes. If two genes *g*_1_ and *g*_2_ are placed together in a common node of the driver tree, we say the pair (*g*_1_, *g*_2_) is in the set of mutual exclusivity relations implied by *T*. Similarly, if the node including *g*_1_ is an ancestor of the node including *g*_2_, we say (*g*_1_, *g*_2_) is in the set of progression relations implied by *T*. For the synthetic data experiments, where the generative progression model is known, we can measure our success in identifying the mutual exclusivity and progression relations. Let *F*_ME_ be our F-score, i.e., the harmonic mean of our precision and recall, for identifying the mutual exclusivity relations. Similarly, let *F*_PR_ be our F-score for identifying the progression relations. We define our overall score, denoted by *F*_overall_, as the harmonic mean of *F*_ME_ and *F*_PR_, i.e.,

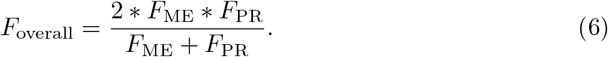

Note that *F*_overall_ ∈ [0, 1] and *F*_overall_ = 1 implies perfect identification of the generative model, while the star tree (which is our initial state) has a *F*_overall_ = 0, as its recall is zero for both mutual exclusivity and progression relations.

Fig. 5 shows the F-scores achieved by our method. As shown in this figure, our inference algorithm finds the exact generative model, or a very similar one for all the cases with error probabilities up to 0.05 and at least 100 tumors in the dataset. The main reason leading to the seemingly poor results for the cases with *ϵ* = 0.1 is that the star tree, which is our initial state, provides likelihood values almost as good as the generative models for such cases. We have provided extensive analysis of the synthetic experiments in S2 Text. This document includes detailed explanations on our evaluation metrics, the precision and recall values, the performance in terms of likelihoods, the inferred error values, and more analysis of the synthetic data experiments.

**Fig 5.**
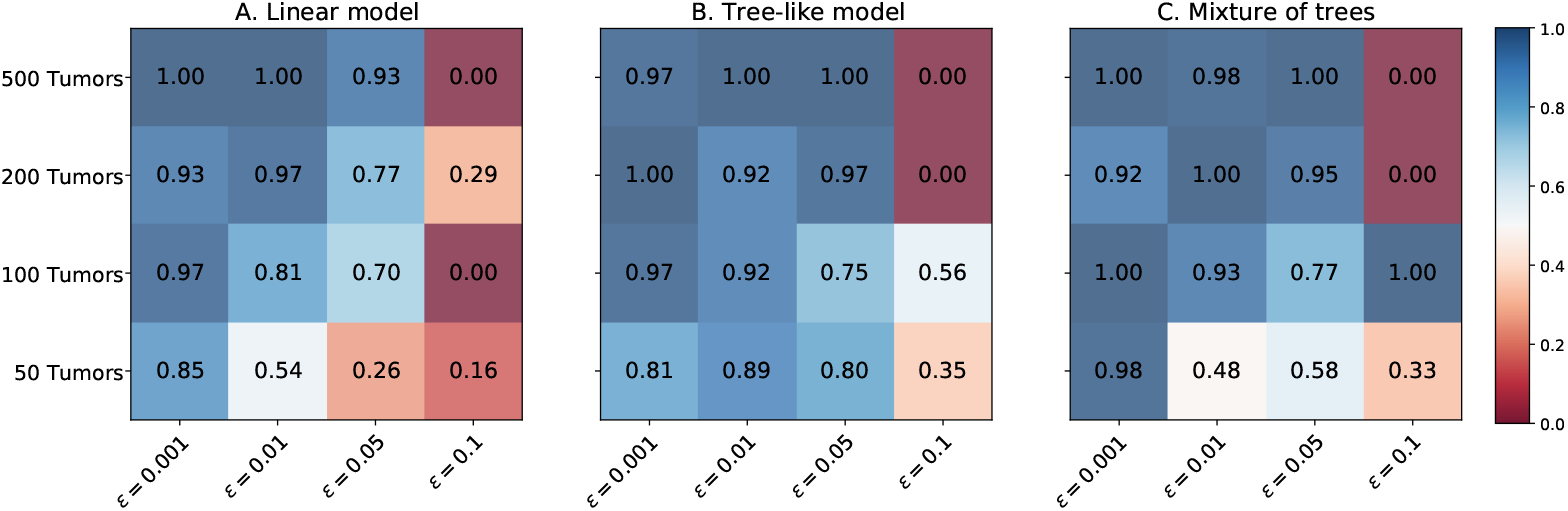
The F-scores achieved in the synthetic data experiments in the case of **A**. Linear model **B**. Tree-like model **C**. Mixture of trees

## 3 Biological Data Analysis

In this section, we present our analysis of several biological datasets. We downloaded mutation-called TCGA data from the GDAC firehose. As the first preprocessing step, we filtered out all silent mutations. We used the genes that meet both following criteria as our “potentially driver genes” to include in the input matrix:

- The gene should be included in the IntoGen’s list of driver genes [11],
- The gene should be highly mutated in the data, with mutations in at least 2 percent of the tumors.

### 3.1 Evaluation metrics

To evaluate how well our resulting progression models explain the datasets, we introduce the following mutual exclusivity and progression scores. These scores provide us with a measure of the support for mutual exclusivity or progression relations among two genes. We use the averaged mutual exclusivity scores of the pairs of genes in each node, or the pairs of genes on two sides of each edge, to assess the performance of our method on the biological datasets.

#### 3.1.1 Mutual exclusivity score

To measure the extent of mutual exclusivity among mutations in a pair of genes (*g*_1_, *g*_2_) placed together in a node, given a dataset *B* of shape *M* ×*N*, we introduce a metric called ME-score. Let 𝒳(*g*) and 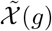 be the number and frequency of mutations in gene *g*, i.e.,

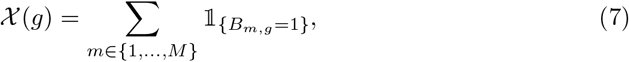

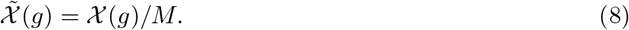

Similarly, we denote the number and frequency of mutations in both *g*_1_ and *g*_2_ by 𝒳 (*g*_1_, *g*_2_) and 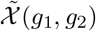, respectively. We define the ME-score of (*g*_1_, *g*_2_) as

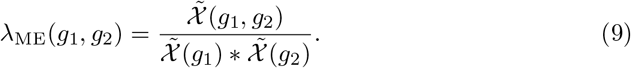

Note that ME-score being zero implies perfect mutual exclusivity among the two genes, and ME-score being one implies the genes’ independence. For the case of *λ*_ME_(*g*_1_, *g*_2_) *<* 1, we wish to measure the statistical significance of the mutual exclusivity signal. To this end, we use a null hypothesis of the genes being independent. Without loss of generality, suppose 𝒳(*g*_1_) ≤ 𝒳(*g*_2_). The p-value will be

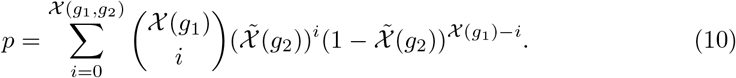

Similarly, when *λ*_ME_(*g*_1_, *g*_2_) *>* 1, we can define a p-value for the mutual inclusivity signal as:

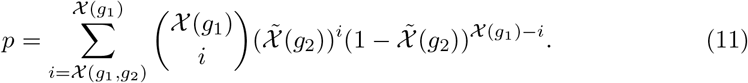

#### 3.1.2 Progression test

To measure the significance of the progression relationship along each edge of the driver tree, we define a metric called PR-score. Consider edge (*u, v*) of the progression model. Let 𝒳_*v*_ and 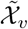 be the number and frequency of tumors having at least one mutation in the genes assigned to node *v*, i.e.,

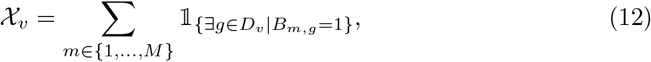

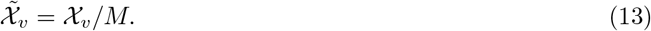

Let *Ƶ* _*v*_ and 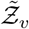 be the number and frequency of tumors mutated in *v*, among the set of tumors with no mutations in the parent node *u*, i.e.,

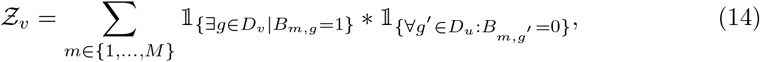

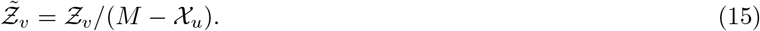

We define PR-score of edge (*u, v*) as

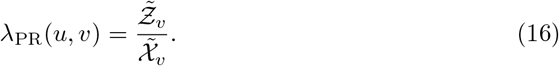

Note that PR-score being zero means that when the parent node is not mutated, there is no mutation in the child, which implies a perfect progression relation. On the other side of the spectrum, a PR-score equal to one implies independence between nodes on two sides of the edge.

For the case of *λ*_PR_(*u, v*) *<* 1, we can calculate a p-value with the null hypothesis of the nodes *u* and *v* being independent as:

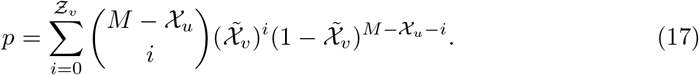

### 3.2 Inferred progression models

In this section, we present our results on the biological datasets. Fig. 6 shows our progression models for three cancer types.

**Fig 6.**
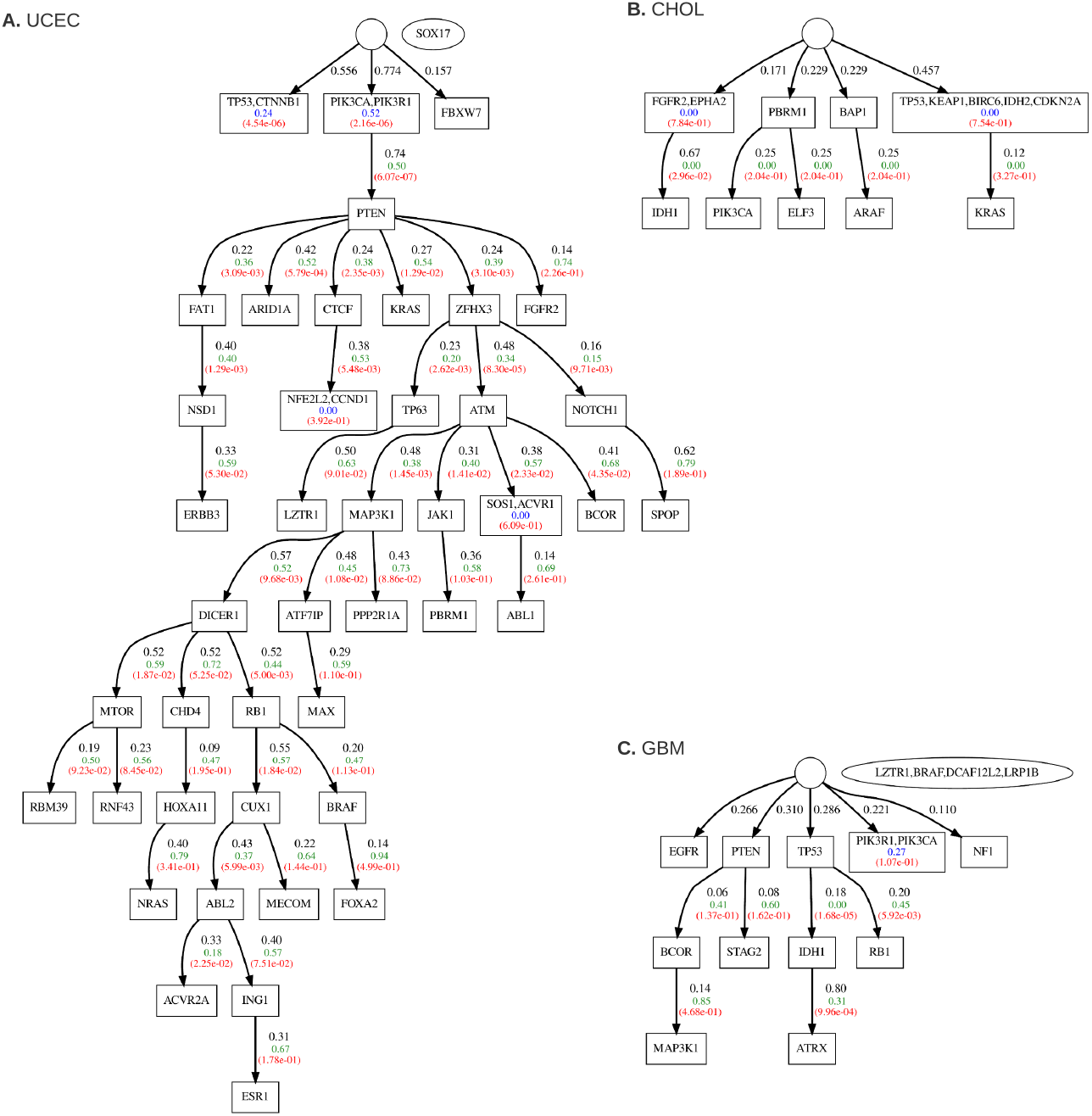
Inferred progression model for A: Uterine corpus endometrial carcinoma (UCEC), B: Cholangiocarcinoma (CHOL), C: Glioblastoma multiforme (GBM). The figure shows the ME-score of the nodes (with at least two genes) in blue. Below the ME-scores, we show the p-values for the mutual exclusivity signals in red. We have the firing probability in black, the PR-score in green, and the corresponding red p-value on each driver tree edge. The oval-shaped node beside the driver tree represents the set of passenger genes if there is any.

#### 3.2.1 Uterine corpus endometrial carcinoma (UCEC)

Our UCEC dataset includes 248 tumors and 48 potentially driver genes. Fig. 6-A shows our progression model for this cancer type. The inferred false positive and false negative probabilities are 0.036 and 0.091, respectively. The likelihood ratio of the resulting model to the star tree is 2.49 * 10^183^, which translates to a per-tumor ratio of 5.49. The inferred model includes several known mutually exclusive oncogenes. The genes TP53 and CTNNB1 have long been known as mutually exclusive oncogenes for liver cancer and hepatocellular carcinoma (HCC) in particular [12]. We have a solid mutual exclusivity signal among these genes here, with a very significant p-value. We have PIK3CA and PIK3R1 from PI3K pathway in the first node of the main branch. PI3K-AKT signaling pathway is known to be one of the most critical pathways involved in cancer [13]. The PTEN tumor suppressor is known to have a high level of interaction with the PI3K signaling pathway [14]. As shown in the figure, the mutations in PI3K precede mutations in PTEN with a very significant progression signal. Our progression model has recovered a broad set of strong progression relations after the PTEN node. Some of these progression signals, such as ZFHX3 preceding ATM, NOTCH1, and TP63, seem interesting and statistically significant enough for further studies.

#### 3.2.2 Cholangiocarcinoma (CHOL)

Our CHOL dataset includes 35 tumors and 14 genes. Fig. 6-B shows our cholangiocarcinoma progression model. As this progression model can construct the exact dataset with no need for false positives or false negatives, the inferred error parameters are zero. The likelihood ratio of the resulting model to the star tree is 2.16 * 10^9^, which translates to a per-tumor ratio of 1.85. While the small number of tumors does not allow for significant mutual exclusivity or progression signals, our inference algorithm has found several interesting perfect mutual exclusivity and progression relations.

#### 3.2.3 Glioblastoma multiforme (GBM)

Our GBM dataset includes 290 tumors and 16 genes. The resulting progression model is shown in Fig. 6-C. Our inferred false positive and false negative probabilities are 0.019 and 0.041, respectively. The likelihood ratio of the resulting model to the star tree is 2.80 * 10^25^, corresponding to a per-tumor likelihood ratio of 1.22. Like the UCEC progression model, we have PIK3CA and PIK3R1, from the PI3K pathway, together in a first-layer node. The tumor suppressors TP53 and PTEN form the first level of the main branches. We have identified solid progression signals from TP53 to IDH1 and RB1 and from IDH1 to ATRX, which suggest further studies on their interrelations.

## Discussion

We have introduced a new model for cancer progression. Our model puts cancer driver genes into sets of mutually exclusive genes and orders them in a tree-like structure, based on the temporal order of driver mutations in them. We introduced an efficient algorithm to compute the likelihood of observing different sets of mutations in each tumor. Using this likelihood calculation procedure within an MCMC algorithm, we find a progression model from cross-sectional data, including a set of mutations observed in a cohort of tumors. After evaluating our inference performance using a comprehensive collection of synthetic data experiments, we used our method to analyze several biological datasets. We introduced two independent evaluation metrics for mutual exclusivity and progression signals among the oncogenes and used them to validate our results. Our inferred progression models for uterine corpus cancer, cholangiocarcinoma, and glioblastoma recovers several well-known mutual exclusivity and progression pattern among specific genes and suggest a broad set of new such relations among the oncogenes.

## Supporting Information

**S1 Text. Method details**.

**S2 Text. Synthetic data experiments**.

## Acknowledgments

This project has received funding from the European Union’s Horizon 2020 research and innovation programme under the Marie Skłodowska-Curie grant agreement MSCA-ITN-2017-766030 and from the Swedish Foundation for Strategic Research grant BD15-0043.

## Notes

### Competing Interest Statement

The authors have declared no competing interest.

