## supplementary text 1 for "Tree of Mutually Exclusive Oncogenes"

### Method details

In this document, we first explain the details of the ToMExO building blocks. In section 1, we present our dynamic programming algorithm for estimating the error parameters. This is the procedure used in the iterations of our inference algorithm. In section 2, we provide the details of a gradient descent approach for estimating the error parameters. We use this procedure to fine-tune our error parameters after deciding on the progression model. In section 3, we provide detailed explanations on our structural moves. Finally, in section 4, we explain our inference algorithm in detail.

We conclude this document with section 4 by explaining our inference algorithm in depth.

#### 1 Empirical Error Estimation

Given the dataset $B$ and a progression model $T = \big(V, E, \{f_v\}_{v\in V}, \{D_v\}_{v\in V}, P\big)$, we want to calculate an estimation for the error parameters $\epsilon$ and $\delta$. We denote the minimum number of false positives and false negatives in $B_{m,v}$ in case of $s_{m,v} = s \in \{0,1\}$ as $Q_{m,v,s} = (n^{\text{FP}}_{m,v,s}, n^{\text{FN}}_{m,v,s})$. We have

$$Q_{m,v,0} = (o_{m,v}, 0), \tag{1}$$

$$Q_{m,v,1} = \begin{cases} (0,1) & , o_{m,v} = 0 \\ (o_{m,v} - 1, 0) & , o_{m,v} \geq 1 \end{cases}. \tag{2}$$

Denoting the minimum number of false positives and false negatives in $B_{m,v\downarrow}$, given $s_{m,v} = s$ as $W_{m,v,s}$, we have

$$W_{m,v,0} = Q_{m,v,0} + \sum_{c\in\mathcal{C}(v)} W_{m,c,0}, \tag{3}$$

$$W_{m,v,1} = Q_{m,v,1} + \sum_{c\in\mathcal{C}(v)} \min_{\|W\|_1}\{W_{m,c,0}, W_{m,c,1}\}. \tag{4}$$

We can calculate these $W$ variables traversing the tree in a post-order manner. Our empirical estimation of the number of false positives and false negatives in the dataset will be:

$$\Big(n_{FP}, n_{FN}\Big) = \sum_{m\in 1,\dots,M} \big(W_{m,\text{root},1} + (o_{m,P}, 0)\big). \tag{5}$$

Denoting the total number of ones and zeros in the dataset by $\mathcal{O}$ and $\mathcal{Z}$, respectively, we have:

$$\Big(\hat{\epsilon}, \hat{\delta}\Big) = \left(\frac{n_{FP}}{\mathcal{Z} - n_{FN} + n_{FP}}, \frac{n_{FN}}{\mathcal{O} - n_{FP} + n_{FN}}\right). \tag{6}$$

Note that only a single post-order traversal over the tree is needed to estimate the error parameters. A pseudo-code for this procedure is provided in Algorithm 1.

**Algorithm 1.** Empirical error estimation

**Input**: $B$, $T = \left(V, E, \{f_v\}_{v \in V}, \{D_v\}_{v \in V}, P\right)$

**Output**: $\hat{\epsilon}, \hat{\delta}$

**for all** $m \in \{1, \ldots, M\}$ **do**
  **for** $v \in$ post-order$(V)$ **do**
    $o_{m,v} = \|B_{m,D_v}\|_1$
    $Q_{m,v,0} = (o_{m,v}, 0)$
    **if** $o_{m,v} = 0$ **then**
      $Q_{m,v,1} = (0, 1)$
    **else**
      $Q_{m,v,1} = (o_{m,v} - 1, 0)$
    $W_{m,v,0} = Q_{m,v,0} + \sum_{c \in \mathcal{C}(v)} W_{m,c,0}$
    $W_{m,v,1} = Q_{m,v,1} + \sum_{c \in \mathcal{C}(v)} \min_{\|W\|_1} \{W_{m,c,0}, W_{m,c,1}\}$
$\left(n_{FP}, n_{FN}\right) = \sum_{m \in 1,\ldots,M} \left(W_{m,\text{root},1} + (\|B_{m,P}\|_1, 0)\right)$
$\left(\hat{\epsilon}, \hat{\delta}\right) = \left(\frac{n_{FP}}{\|1-B\|_1 - n_{FN} + n_{FP}}, \frac{n_{FN}}{\|B\|_1 - n_{FP} + n_{FN}}\right)$

### 2 Gradient Descent for Error Estimation

As another approach for error parameter estimation, we can use Gradient Descent to find an at least locally optimal estimate of the parameters $\epsilon$ and $\delta$, given a progression model $T$ and the dataset $B$. Instead of working with the likelihood itself, our objective is to maximize the log-likelihood, which can be written as

$$l(B|T, \epsilon, \delta) = \sum_{m \in \{1,\ldots,M\}} \log(p(B_{m,:}|T, \epsilon, \delta)). \tag{7}$$

We have

$$\frac{\partial l(B|T, \epsilon, \delta)}{\partial \epsilon} = \sum_{m \in \{1,\ldots,M\}} \frac{\partial \log p(B_{m,:}|T, \epsilon, \delta)}{\partial \epsilon}, \tag{8}$$

where

$$\frac{\partial \log p(B_{m,:}|T, \epsilon, \delta)}{\partial \epsilon} = \frac{\partial \log \Psi_{m,\text{root}}}{\partial \epsilon} + \frac{o_{m,P}}{\epsilon} - \frac{z_{m,P}}{1-\epsilon}. \tag{9}$$

We can compute the derivatives of the $\Psi$ variables at the root node with a single post-order traversal of the driver tree. Following the formula used for likelihood calculations, we have

$$\frac{\partial \log \Lambda_{m,v}}{\partial \epsilon} = \frac{1}{\Lambda_{m,v}}\Bigg(\frac{o_{m,v}}{o_{m,v}+z_{m,v}}(1-\delta)\epsilon^{o_{m,v}-2}(1-\epsilon)^{z_{m,v}-1}\big(\epsilon(1-o_{m,v}-z_{m,v})+o_{m,v}-1\big)$$

$$+\frac{z_{m,v}}{o_{m,v}+z_{m,v}}\delta\epsilon^{o_{m,v}-1}(1-\epsilon)^{z_{m,v}-2}\big(o_{m,v}-\epsilon(o_{m,v}+z_{m,v}-1)\big)\Bigg) \tag{10}$$

$$\frac{\partial \log \Omega_{m,v}}{\partial \epsilon} = \frac{o_{m,v}}{\epsilon} - \frac{z_{m,v}}{1-\epsilon} + \sum_{c\in\mathcal{C}(v)} \frac{\partial \log \Omega_{m,v}}{\partial \epsilon} \tag{11}$$

$$\frac{\partial \log \Psi_{m,v}}{\partial \epsilon} = \frac{\partial \log \Lambda_{m,v}}{\partial \epsilon} + \sum_{c\in\mathcal{C}(v)} \frac{f_c\Psi_{m,c}\frac{\partial \log \Psi_{m,c}}{\partial \epsilon} + (1-f_c)\Omega_{m,c}\frac{\partial \log \Omega_{m,c}}{\partial \epsilon}}{f_c\Psi_{m,c}+(1-f_c)\Omega_{m,c}} \tag{12}$$

Similarly, for the $\delta$ variable we have

$$\frac{\partial l(B|T,\epsilon,\delta)}{\partial \delta} = \sum_{m\in\{1,\ldots,M\}} \frac{\partial \log p(B_{m,:}|T,\epsilon,\delta)}{\partial \delta}, \tag{13}$$

where

$$\frac{\partial \log p(B_{m,:}|T,\epsilon,\delta)}{\partial \delta} = \frac{\partial \log \Psi_{m,\text{root}}}{\partial \delta}. \tag{14}$$

To compute the derivatives of the $\Psi$ variables, we have

$$\frac{\partial \log \Lambda_{m,v}}{\partial \delta} = \frac{1}{\Lambda_{m,v}}\left(\frac{\epsilon^{o_{m,v}-1}(1-\epsilon)^{z_{m,v}-1}\big(\epsilon(z_{m,v}+o_{m,v})-o_{m,v}\big)}{z_{m,v}+o_{m,v}}\right) \tag{15}$$

$$\frac{\partial \log \Omega_{m,v}}{\partial \delta} = \sum_{c\in\mathcal{C}(v)} \frac{\partial \log \Omega_{m,v}}{\partial \delta} \tag{16}$$

$$\frac{\partial \log \Psi_{m,v}}{\partial \delta} = \frac{\partial \log \Lambda_{m,v}}{\partial \delta} + \sum_{c\in\mathcal{C}(v)} \frac{f_c\Psi_{m,c}\frac{\partial \log \Psi_{m,c}}{\partial \delta} + (1-f_c)\Omega_{m,c}\frac{\partial \log \Omega_{m,c}}{\partial \delta}}{f_c\Psi_{m,c}+(1-f_c)\Omega_{m,c}} \tag{17}$$

Having the derivatives, we can use any gradient-based optimization method to find $\epsilon$ and $\delta$.

Note that while we can compute the gradients using a single traverse of the driver tree, the gradient descent methods are iterative processes themselves. Hence, this approach's computational complexity is much higher than the procedure that obtains the empirical error estimates. As the overall performance of our method does not degrade much by using the empirical error estimation, we use the gradient descent algorithm only for fine-tuning the error parameters $\epsilon$ and $\delta$ after finding the final progression model.

### 3 Structural Moves

To sample progression models using a Metropolis-Hasting approach, we need to have structural moves involving changes in the topology of the driver tree and the assignment of the genes to the nodes. We have designed 14 types of moves to propose new progression models, which may then be accepted based on the Metropolis-Hasting acceptance criteria.

The moves are introduced one by one in the following. Note that whenever we need to *select* one or a set of nodes or genes, we do it uniformly from the set of candidate choices. As an illustrative example, let the progression model shown in Fig. 1-A be the model in hand, i.e., the current state of the chain. Fig. 1-B to O show example proposed models for each one of our move types.

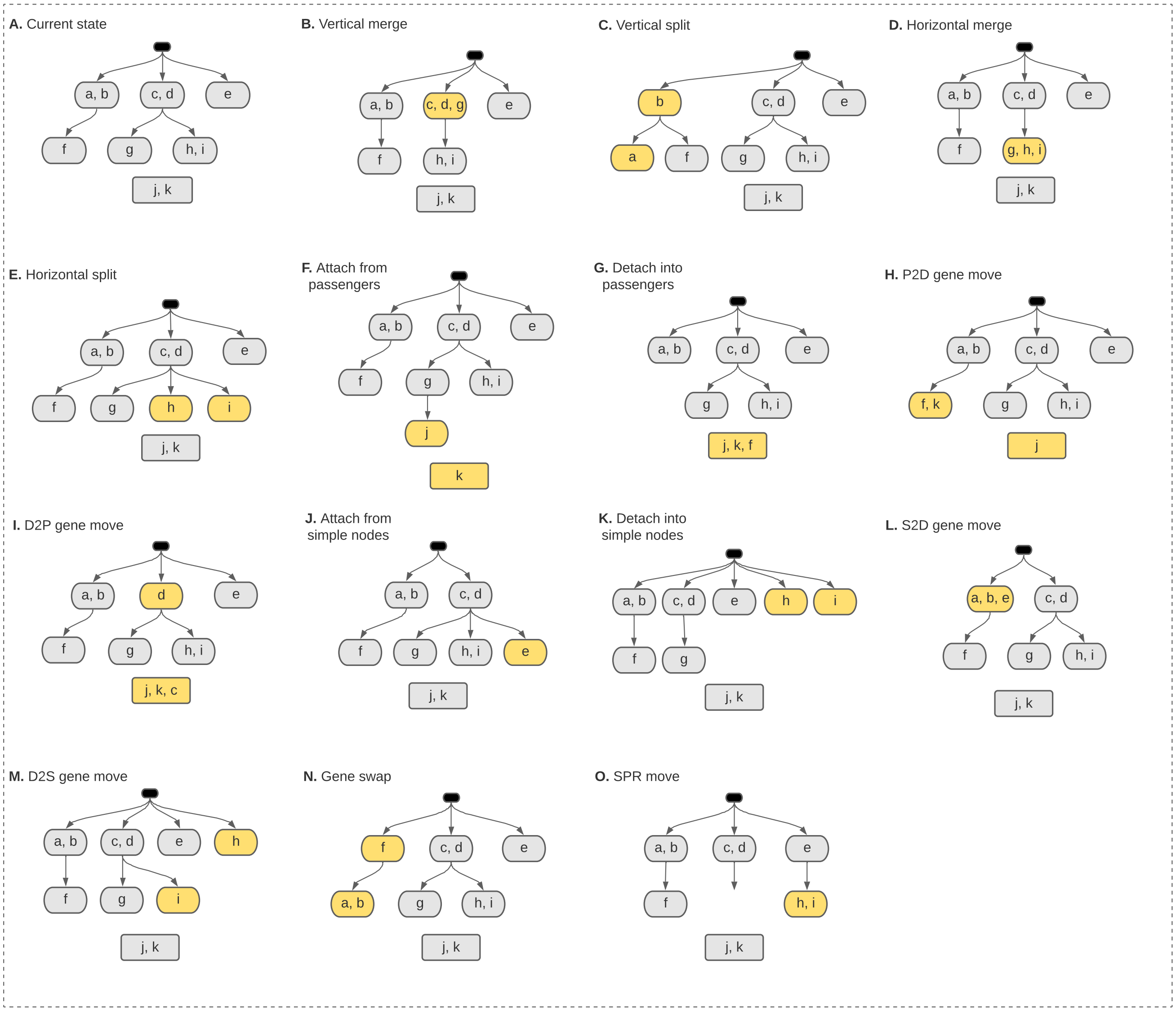


Figure 1: Different types of structural moves used in our MCMC inference algorithm.

#### 3.1 Vertical merge

For this move, we merge a leaf node into its parent. We select the leaf node from the leaves that are not in the first layer of the tree so that the resulting tree has a standard structure with no genes in the root node. Fig. 1-B shows an example of such moves.

#### 3.2 Vertical split

For this move, we choose a node with at least two genes, select a proper non-empty subset of its genes and move them into a new child node. Fig. 1-C shows an example of such moves.

#### 3.3 Horizontal merge

For this move, we merge a pair of sibling leaves together. Fig. 1-D shows an example of such moves.

#### 3.4 Horizontal split

For this move, we split a leaf node into two sibling leaves. To this end, we select a leaf node that includes at least two genes. We partition the selected node's genes into two non-empty subsets and put them into the resulting sibling nodes. Fig. 1-E shows an example of such moves.

#### 3.5 Attach from Passengers

For this move, we select a non-empty subset of the passenger genes, form a new node with them, and select its parent node from the existing nodes of the tree. Fig. 1-F shows an example of such moves.

#### 3.6 Detach into Passengers

For this move, we select a leaf node, remove it and put its genes into the set of passengers. Fig. 1-G shows an example of such moves.

#### 3.7 P2D gene move

This is the passenger-to-driver gene move. For this move, we select a gene from the set of passengers and move it to one of the nodes in the driver tree. Fig. 1-H shows an example of such moves.

#### 3.8 D2P gene move

This is the driver-to-passenger gene move. For this move, we select a node with at least two genes in it, pick one of its genes, and move it to the set of passengers. Fig. 1-I shows an example of such moves.

#### 3.9 Attach from simple nodes

With this move, we select a non-empty subset of simple nodes to remove and put their genes in a new node. We attach this new node to a parent node, which

- is not among the selected subset of simple nodes,
- is not the root, if the selected subset includes one or two simple nodes (as these cases will correspond to "no move" and "horizontal merge", respectively).

Fig. 1-J shows an example of such moves.

#### 3.10 Detach into simple nodes

With this move, we remove a leaf node and put its genes into new simple nodes. We choose a leaf node, which

- is not a simple node, as it would lead to no move
- is not a first layer node with exactly two genes, as this case is covered in "horizontal split".

Fig. 1-K shows an example of such moves.

##### 3.11 S2D gene move

This is the simple-to-driver gene move. For this move, we select a simple node, remove it and put its genes in a node, which

- is not the root node,
- is not a first-level node with no children (to not conflict with ”horizontal merge”).

Fig. 1-L shows an example of such moves.

##### 3.12 D2S gene move

This is the driver-to-simple gene move. For this move, we select a node in the driver tree, which

- has at least two genes
- is not a first-level node with no children (D2S type of moves for such nodes are covered in ”horizontal split”).

We then select one of the genes in the selected node and move it to a new first-level node, forming a simple node. Fig. 1-M shows an example of such moves.

##### 3.13 Gene swap

For this move, we select a node and swap its genes with those of its parent. We choose our node from those not in the first layer, as the parent node should not be the empty root node. Fig. 1-N shows an example of such moves. It is worth mentioning that the forward and backward probabilities are equal for this move, making it easier to compute the acceptance ratio.

##### 3.14 SPR move

This is the well-known subtree pruning and regrafting move. For this move, we select a node, which

- is not the root,
- is not a simple node.

We then select a new parent for the selected node that

- is not in the selected node's subtree (including the selected node itself),
- is not the root node, if the selected node includes one gene and has no children (as it would result in a new simple node).

Note that the simple nodes are the single-gene leaves in the first layer of the tree. As the “attach from simple nodes” covers the SPR moves that relocate the simple nodes, we do not move them here. Note that the SPR move has equal forward and backward probabilities, which is useful for calculating the acceptance ratio. Fig. 1-O shows an example of such moves.

**Algorithm 2.** Likelihood calculation

**Input**: $B$, $T = \big(V, E, \{f_v\}_{v \in V}, \{D_v\}_{v \in V}, P\big)$, $\epsilon$, $\delta$
**Output**: $p(B|T, \epsilon, \delta)$
**for all** $m \in \{1, \dots, M\}$ **do** ▷ Calculate $p(B_{m,:}|T, \epsilon, \delta)$
  **for** $v \in$ post-order$(V)$ **do**
    $o_{m,v} = \|B_{m,D_v}\|_1$
    $z_{m,v} = \|1 - B_{m,D_v}\|_1$
    $\Lambda_{m,v} = \frac{o_{m,v}}{o_{m,v}+z_{m,v}}(1-\delta)\epsilon^{o_{m,v}-1}(1-\epsilon)^{z_{m,v}} + \frac{z_{m,v}}{o_{m,v}+z_{m,v}}\delta\epsilon^{o_{m,v}}(1-\epsilon)^{z_{m,v}-1}$
    $\Gamma_{m,v} = \epsilon^{o_{m,v}}(1-\epsilon)^{z_{m,v}}$
    $\Psi_{m,v} = \Lambda_v \prod_{c \in \mathcal{C}(v)} \big(f_c \Psi_c + (1 - f_c)\Omega_c\big)$
    $\Omega_{m,v} = \Gamma_v \prod_{c \in \mathcal{C}(v)} \Omega_c$
  $p(B_{m,:}|T, \epsilon, \delta) = \Psi_{m,\text{root}} \epsilon^{\|B_{m,P}\|_1}(1-\epsilon)^{\|1-B_{m,P}\|_1}$
$p(B|T, \epsilon, \delta) = \prod_{m \in \{1,\dots,M\}} p(B_{m,:}|T, \epsilon, \delta)$

### 4 Inference Algorithm

Building on our dynamic programming procedure for likelihood calculation (see Algorithm 2), various MCMC approaches can be used for inference. We use the algorithm described in Algorithm 3, where the firing probability of each edge $(u, v)$, denoted by $f_v$ is set to a specific empirical estimation, as

$$f_v = \mathcal{Y}_{B,v} / \mathcal{X}_{B,u}, \tag{18}$$

where

$$\mathcal{X}_{B,u} = \sum_{m \in \{1,\dots,M\}} \mathbb{1}_{\{\exists g \in D_u | B_{m,g}=1\}}, \tag{19}$$

$$\mathcal{Y}_{B,v} = \sum_{m \in \{1,\dots,M\}} \mathbb{1}_{\{\exists g \in D_v | B_{m,g}=1\}} * \mathbb{1}_{\{\exists g' \in D_u | B_{m,g'}=1\}}. \tag{20}$$

Note that $\mathcal{X}_{B,u}$ estimates the number of tumors with a mutation in $u$ (the parent node), and $\mathcal{Y}_{B,v}$ estimates the number of tumors with mutations in both $v$ and $u$. Hence, $f_v = \mathcal{Y}_{B,v}/\mathcal{X}_{B,u}$ can be seen as an empirical estimation of $f_v$.

In our inference algorithm, we use our empirical error estimation (Algorithm 1) instead of sampling the error parameters $\epsilon$ and $\delta$. We use a Metropolis-Hasting framework, where in each iteration we choose a move type and generate a proposal progression model accordingly. We then use Algorithm 1 to calculate the corresponding error parameters $\epsilon$ and $\delta$, which form the proposed sample together with the progression model. The proposed sample may then get accepted as the next state of the chain with a certain probability. Let $T^c_{i-1}$, $\epsilon^c_{i-1}$ and $\delta^c_{i-1}$ denote the progression model and error parameters in chain $c$ and iteration $i-1$. Denoting the proposed sample by $(\hat{T}^c_i, \hat{\epsilon}^c_i, \hat{\delta}^c_i)$, the acceptance probability is calculated as

$$p^c_{i,\text{accept}} = \frac{p(B|\hat{T}^c_i, \hat{\epsilon}^c_i, \hat{\delta}^c_i)}{p(B|T^c_{i-1}, \epsilon^c_{i-1}, \delta^c_{i-1})} * \frac{p^c_{i,\text{backward}}}{p^c_{i,\text{forward}}}, \tag{21}$$

where $p^c_{i,\text{forward}}$ and $p^c_{i,\text{backward}}$ are the forward and backward probabilities.

At the end of the iterations, we pick the sample with the maximum likelihood and fine-tune the parameters $\epsilon$ and $\delta$ using gradient descent, as detailed in section 2. A pseudo-code of our inference algorithm is provided in Algorithm 3.

**Algorithm 3.** ToMExO inference algorithm

**Input**: $B$, $n_{\text{chains}}$, $n_{\text{iterations}}$
**Output**: $T = \big(V, E, \{f_v\}_{v\in V}, \{D_v\}_{v\in V}, P\big)$, $\epsilon$, $\delta$

**for all** $c \in \{1, ..., n_{\text{chains}}\}$ **do**
  Initialize $T_0^c$ to a star-tree ▷ Using Eq. (18) for $\{f_v\}_{v\in V(T_0^c)}$
  Calculate $\epsilon_0^c$ and $\delta_0^c$ using Algorithm 1 ▷ Empirical error estimation
  **for all** $i \in \{1, ..., n_{\text{iterations}}\}$ **do**
    Choose a move type uniformly
    Generate a proposal sample $\hat{T}_i^c$
    Calculate $\hat{\epsilon}_i^c$ and $\hat{\delta}_i^c$ using Algorithm 1 ▷ Empirical error estimation
    Calculate acceptance ratio $p_{i,\text{accept}}^c$ using Eq. (21)
    Draw $q \sim \text{Uniform}(0, 1)$
    **if** $q < p_{i,\text{accept}}^c$ **then**
      $T_i^c = \hat{T}_i^c$, $\epsilon_i^c = \hat{\epsilon}_i^c$, $\delta_i^c = \hat{\delta}_i^c$ ▷ Accepted proposal
    **else**
      $T_i^c = T_{i-1}^c$, $\epsilon_i^c = \epsilon_{i-1}^c$, $\delta_i^c = \delta_{i-1}^c$ ▷ Rejected proposal
Choose the best sample $(T, \epsilon, \delta) = \arg\max_{(T_i^c, \epsilon_i^c, \delta_i^c)} p(B|T_i^c, \epsilon_i^c, \delta_i^c)$
Optimize $\epsilon$ and $\delta$ values using gradient descent (section 2)
