## supplementary text 2 for "Tree of Mutually Exclusive Oncogenes"

### Synthetic data experiments

In this document, we provide further details on the results of our synthetic data experiments. As explained in the paper, we have three generative progression models shown in Fig. 4 of the paper. For each case, we constructed 16 datasets with error rates in $\{0.001, 0.01, 0.05, 0.1\}$ and the number of tumors in $\{50, 100, 200, 500\}$. Note that while we use the same values for the probability of false-positive $\epsilon$ and false-negative $\delta$ errors, the inference algorithm views them as independent variables. We used the inference algorithm with 10 chains and $100k$ samples for all the experiments.

Fig. 1 shows the precision and recall values for identifying the mutual exclusivity and progression relations. The resulting F-scores are shown in Fig. 2a and Fig. 2b. Fig. 2c shows our overall scores $F_{\text{overall}}$ for the synthetic data experiments. As shown in these figures, our inference algorithm provides excellent results when the error probabilities are up to 0.05, especially when the number of tumors is 100 or more. For most cases with error probability 0.1, our inference algorithm tends to stick to the star tree or a progression model implying very few progression and mutual exclusivity relations, resulting in very low F-scores. For some other cases, when we have only 50 tumors in the data, some relations in deeper levels of the progression model are difficult to identify, leading to low F-scores. Consider the linear model with $\epsilon = 0.01$, for example. As shown in Fig. 1, when we have only 50 tumors, the precision and recall values are not satisfactory. Notably, the recall values for both progression and mutual exclusivity relations are way lower than their corresponding precision values. The problem in this example case is solved as the number of tumors in the data increases.

To better explain the results, Fig 3a shows the likelihood ratio (per tumor) of the star tree to the generative progression model, i.e.,

$$\left(\frac{p(B|T_0, \epsilon_0, \delta_0)}{p(B|T_g, \epsilon_g, \delta_g)}\right)^{1/M}, \quad (1)$$

where $(T_0, \epsilon_0, \delta_0)$ denotes the star tree and it's corresponding error values, and $(T_g, \epsilon_g, \delta_g)$ are the generative tree and error variables. As shown in the figure, when the error rates increase, the generative model loses its edge over the star tree, and the star tree gets a likelihood very close to the generative model. As a result, this is more difficult for the inference algorithm to improve from the initial state, which is the star tree, as it already provides a very competitive likelihood. Fig 3b shows the likelihood ratio (per tumor) of the output tree to the generative progression model. This figure shows that for almost all the cases with error probabilities up to 0.05, we have found a model with a likelihood at least as good as the generative model. As a sanity check, we can see that in all 48 cases, the output gives a likelihood ratio at least as high as the initial star tree. Interestingly, even for some of the cases where $\epsilon = \delta = 0.1$, we have outputs with higher likelihoods than the generative models. Take the dataset with 200 tumors and $\epsilon = \delta = 0.1$ in the linear model scenario as an example. We have started from a likelihood ratio of 0.88 and reached 1.03. This means that the output has a higher likelihood than the generative model. However, as the output is not exactly equal to the generative model, the overall F-score is as low as 0.29 for this dataset (see Fig. 2c). To further investigate this specific case, the output sample is shown in Fig. 4. The fine-tuned error parameters for this sample are $\epsilon = 0.0531$ and $\delta = 02579$.

The fine-tuned error parameters provided in the outputs are shown in Fig. 5a and Fig. 5b. These figures show that the estimated error parameters are typically lower than the generative values, as expected. The figures also show a general trend of increasing inferred values as the generative error parameter increases. We see that in some seemingly strange cases, the inferred $\epsilon$ and $\delta$ values are very small, while the generative error probability is equal to 0.1. Take the case of the linear model with 500 tumors and $\epsilon = 0.1$, for instance. The reason for such poor performances in estimating the error parameters is that the output trees are the star trees or very similar to them. Having the star tree as the output, we do not need a high error probability to get a good likelihood, mostly because of the perfectly matching firing probabilities.

While we work with independent false positive and false negative probabilities, our model and inference algorithm can easily be modified to work with a single error parameter. As equal $\epsilon$ and $\delta$ parameters are used while constructing our synthetic datasets, one may suggest exploring what would happen if we had tied these parameters together in our inference procedure. Fig. 6 answers this question. Fig. 6a shows the overall F-scores for the outputs of an inference algorithm that works with a single error parameter. Comparing these results with the ones we got from our inference algorithm in Fig. 2, we see that the results are quite similar, with a few slightly better results and a few slightly worse ones. Fig. 6b shows the inferred error parameters. As expected, the estimated error parameters follow the same patterns observed in the case of two independent false positive and false negative error parameters.

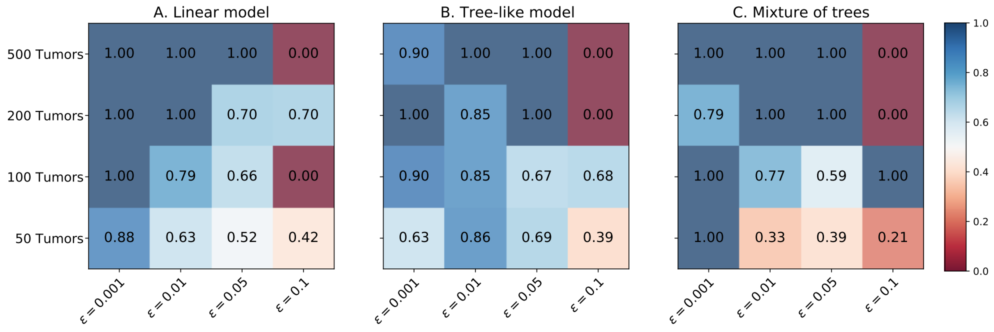


(a) Progression precision

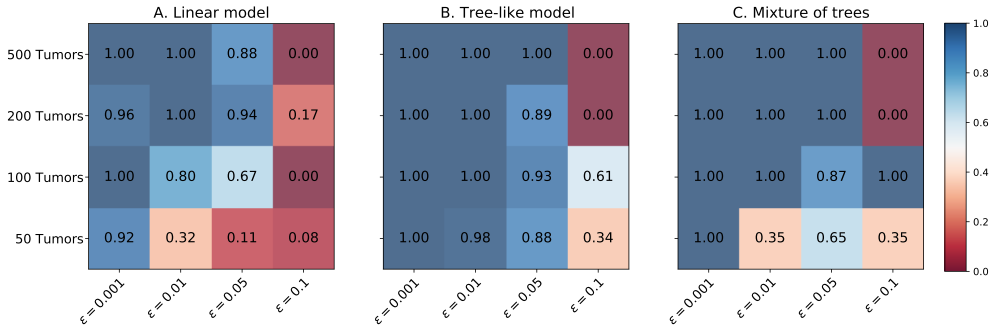


(b) Progression recall

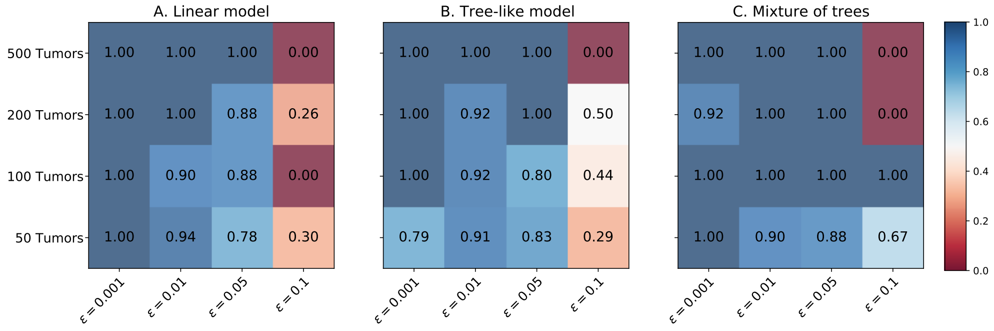


(c) Mutual exclusivity precision

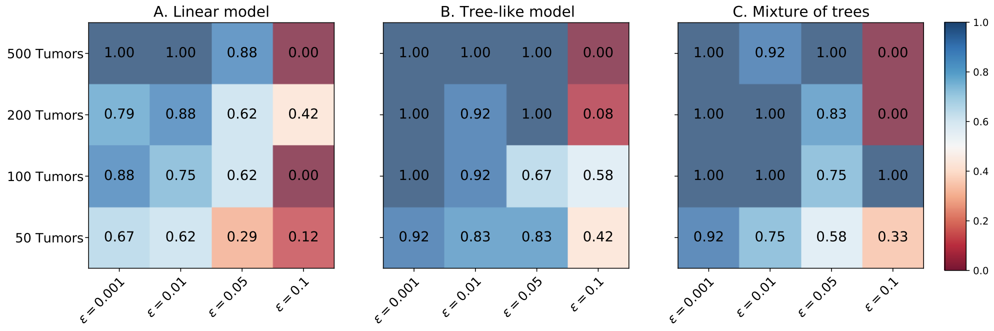


(d) Mutual exclusivity recall

Figure 1: The resulting precision and recall values

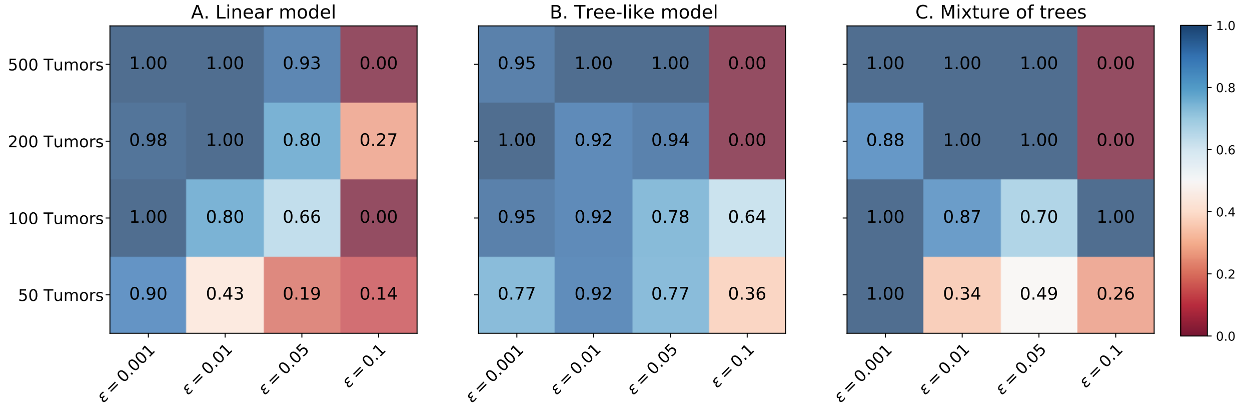


(a) Progression F-scores, $F_{\mathrm{PR}}$

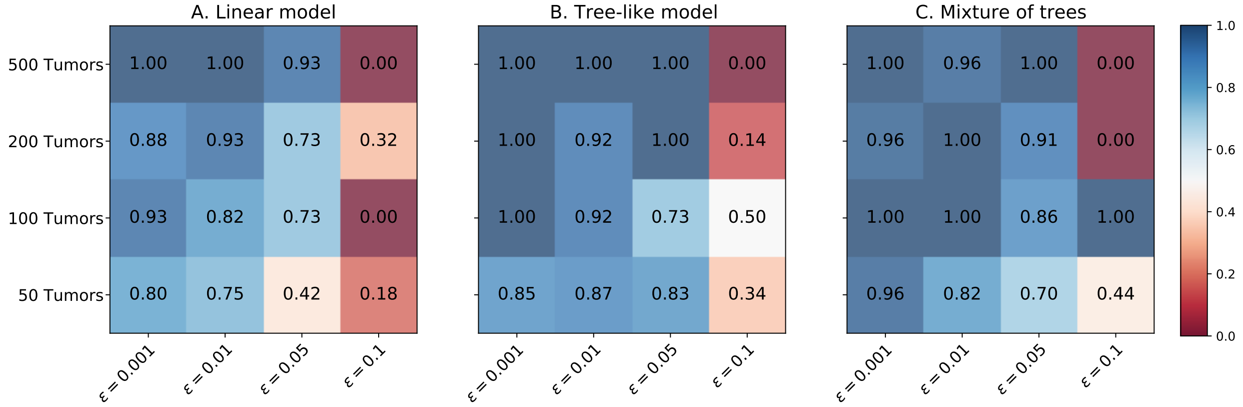


(b) Mutual exclusivity F-scores, $F_{\mathrm{ME}}$

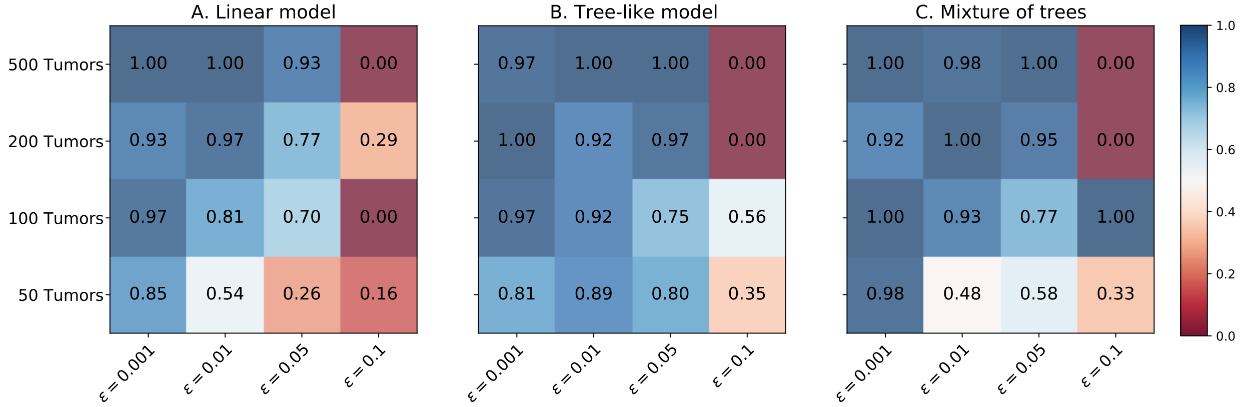


(c) Overall F-scores, $F_{\mathrm{overall}}$

Figure 2: The resulting F-scores

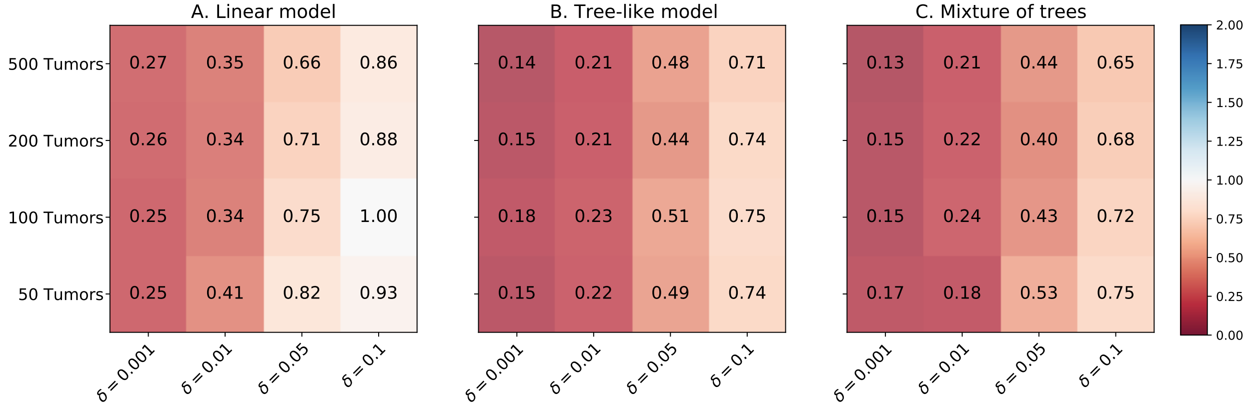


(a) Likelihood ratio (per tumor) of the star tree to the generative model

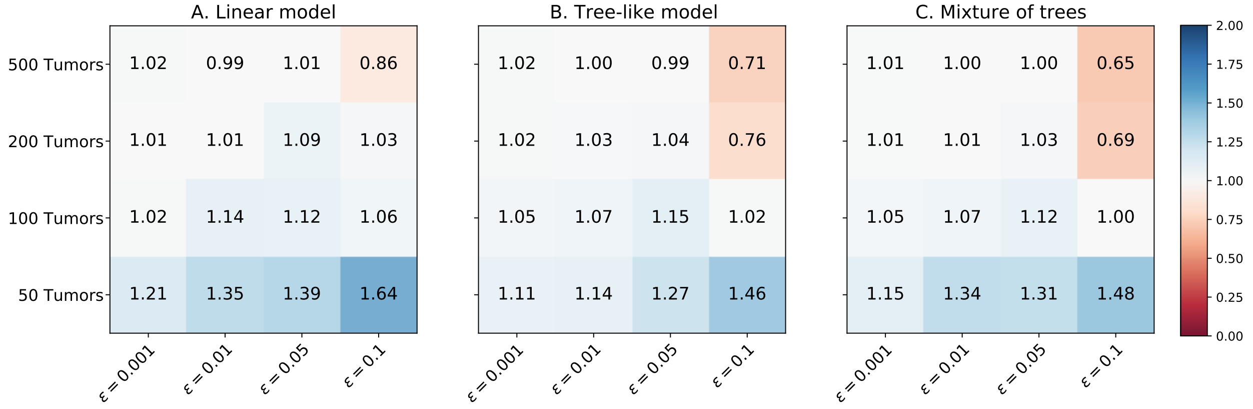


(b) Likelihood ratio (per tumor) of the output tree to the generative model

Figure 3: The likelihood ratios

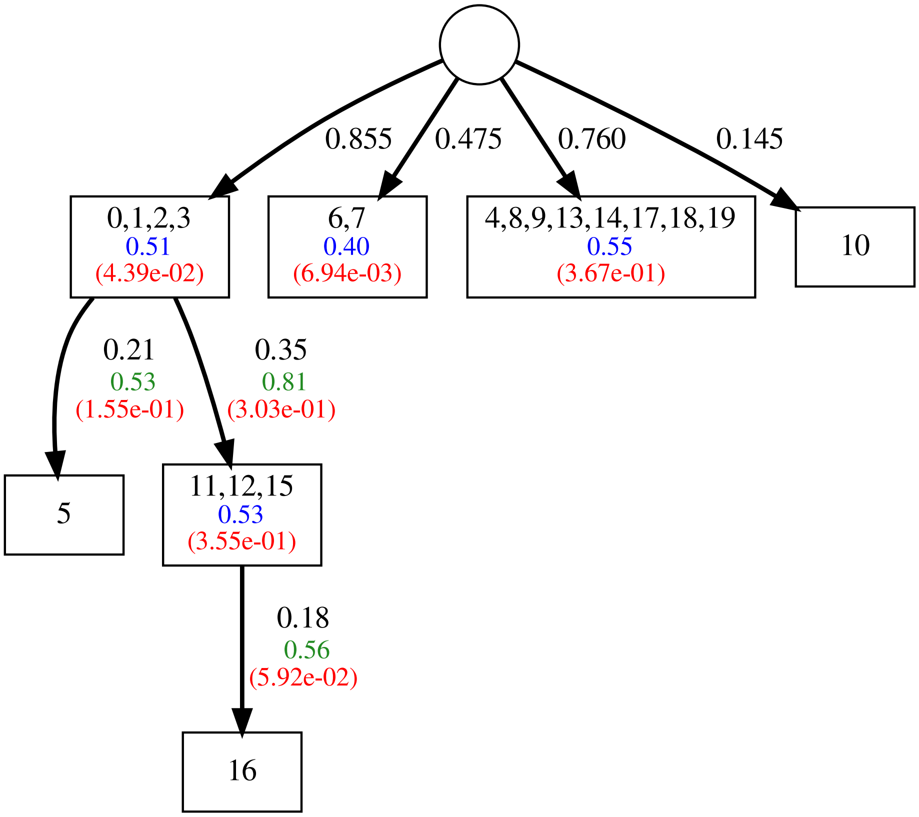


Figure 4: The output tree for the case of linear generative model, 200 tumors and error parameters $\epsilon = \delta = 0.1$. The genes placed in each node are shown by their indices in black. The blue and green numbers show the mutual exclusivity and progression scores, respectively. These scores are method-independent evaluation metrics, between 0 and 1, with 0 corresponding to the perfect case. The red numbers in the figure are the p-values, showing the significance of the mutual exclusivity and progression signals. For details on the mentioned scores and p-values, see section 3.1 of the paper.

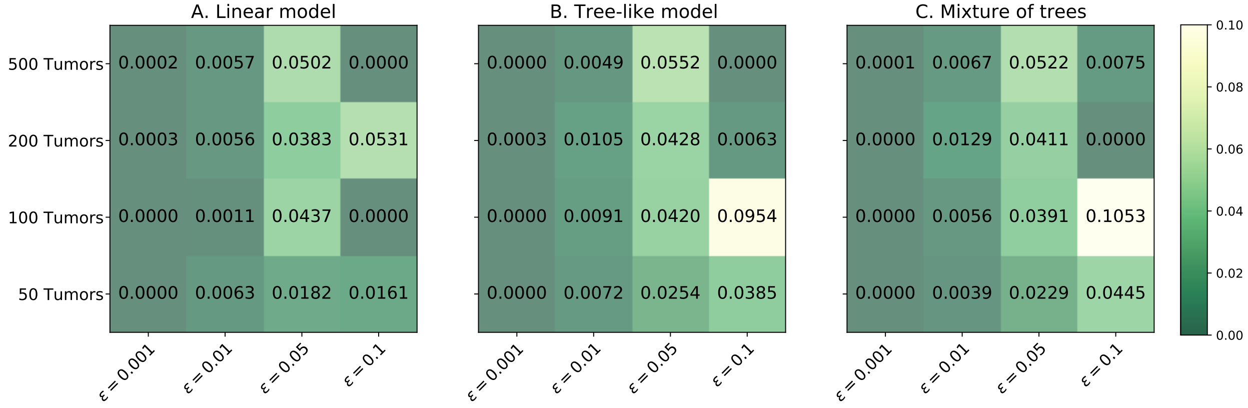


(a) Probability of false positive, $\epsilon$

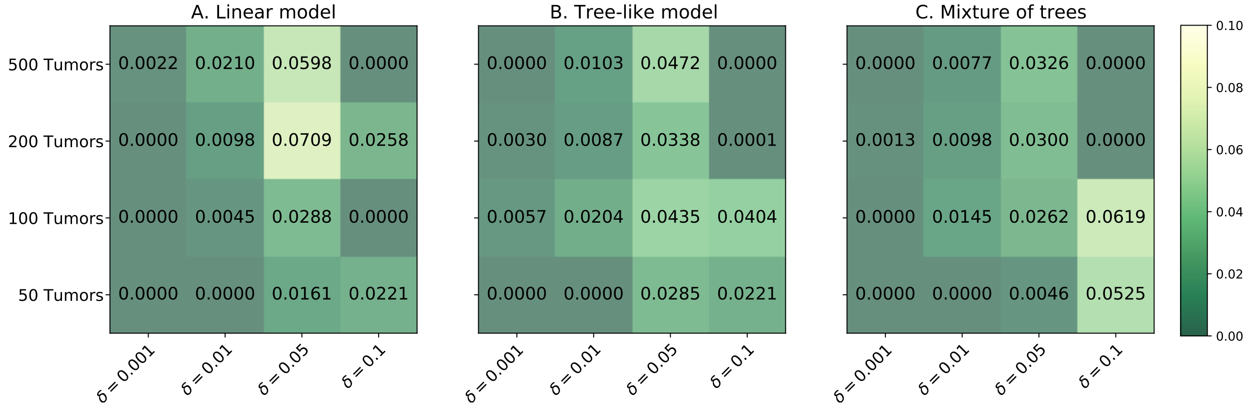


(b) Probability of false negative, $\delta$

Figure 5: The inferred error parameters

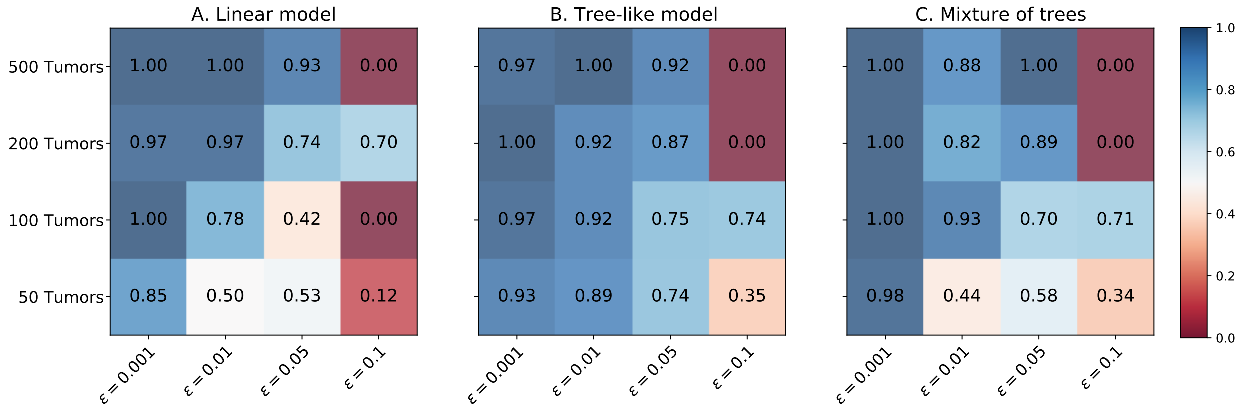


(a) Overall F-scores, $F_{\text{overall}}$ (for the case of a single error parameter)

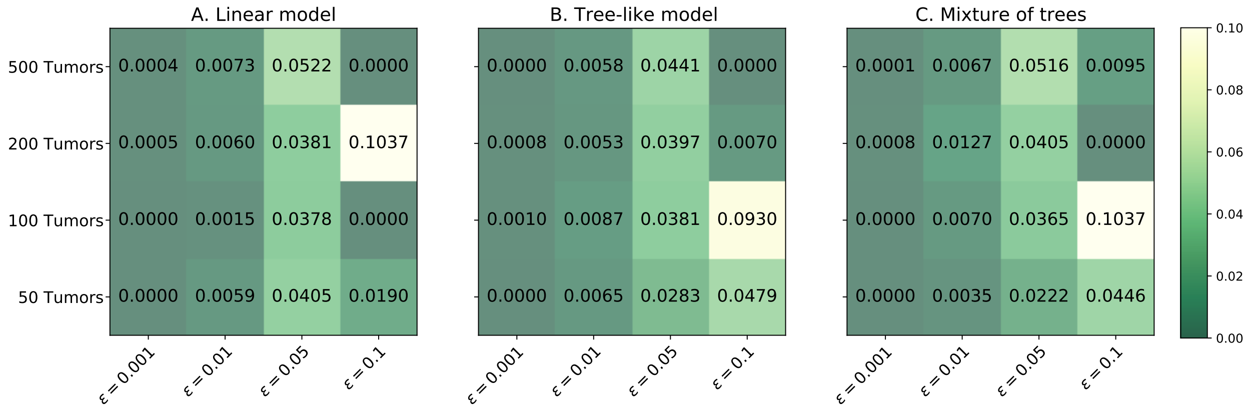


(b) Inferred error probabilities (for the case of a single error parameter)

Figure 6: The results for the case of inference algorithm with one error parameter
